# Macroevolutionary Rates of Species Interactions: Approximate Bayesian Inference from Cophylogenies

**DOI:** 10.1101/2025.05.15.653894

**Authors:** Yichao Zeng, Cristian Román-Palacios

## Abstract

Understanding the macroevolutionary dynamics of species interactions such as parasitisms, commensalisms, and mutualisms is an important goal in evolutionary ecology. To this end, statistical inference from extant cophylogenetic systems holds immense potential. However, such inference cannot yet handle the speciation-extinction dynamics that occur simultaneously in the host and symbiont clades on the same timescale. Here we present an Approximate Bayesian Computation (ABC) approach that, while taking into account host and symbiont extinction, infers rates of four types of speciation from a cophylogenetic system: (i) host speciation, (ii) symbiont speciation without host-switching, (iii) symbiont speciation with host-switching, and (iv) cospeciation. The new ABC approach relies on a novel design of summary statistics combining both size-based (i.e., tree sizes) and size-free summary statistics (i.e., the normalized distribution of Branch Length Differences - BLenD) of the cophylogeny. Convergence analyses show that the combined design of summary statistics outperforms size-based or size-free summary statistics alone – achieving satisfactory accuracy in detecting rate heterogeneity between the four types of speciation. Our ABC approach allows the user to infer the predominant mode of speciation within a given cophylogenetic system. The approach is demonstrated with an application to a cophylogenetic dataset of commensalism, in which beetles of one genus mimic those of another. In this system, we identify host speciation as the predominant process with the fastest rate (in events per unit time) among all four types of speciation (i.e., 4.3-5.1 times faster than the median among all four types of speciation). Understanding how and why different types of speciation may predominate in different cophylogenetic systems can have implications for various areas in ecology and evolution such as host conservatism, trait-driven diversification, pathogen spillover risk, and parasite extinction risk. This new approach highlights the need and potential for future efforts to compile time-calibrated cophylogenies.

## 1 Introduction

Biologists have long sought to understand the relationship between macroevolution and ecological interactions. Early examples include how Darwin explains the diversity of life as a result of natural selection in his seminal book, using examples where natural selection occurs as a result of organisms’ “struggle for life” against their competitors and natural enemies (Darwin, 1859). A recent comprehensive synthesis has consolidated the macroevolution of species interactions as a rapidly-evolving subfield of ecology and evolution with deep historical roots (see Hembry and Weber 2020 for an overview). Several other syntheses have separately examined the effects of species interactions on macroevolution, reaching varying conclusions about whether generalities exist in how species interactions influence clade diversification (Jablonski, 2008; Harmon et al., 2019; Zeng and Wiens, 2021; Kaur and Pennell, 2023). Another body of research has focused on the effects of macroevolution on species interaction networks. Such examples include how macroevolutionary stability shapes the intricate architecture of species interaction networks (Burin et al., 2021) and statistical inference of the evolutionary history of species interaction networks (Braga et al., 2020, 2021). Recent years have also seen significant advances in theoretical work that, by modeling how speciation arises from mutation in a (meta)community with ongoing species interactions, bridges the microevolution-macroevolution gap and provides mechanistic insights into the evolution of ecological communities (Aguilée et al., 2018; Coelho and Rangel, 2018; Maliet et al., 2020; Chaparro-Pedraza et al., 2022; Pontarp et al., 2024; Zeng and Hembry, 2024).

In particular, the joint macroevolutionary history of organisms that interact in pairs (i.e., bipartite interactions) has attracted much attention over the decades. Examples of bipartite interactions include those between plants and pollinators, plants and seed dispersers, hosts and parasites, and between plants and herbivores (see Bronstein 2015 for an overview). Coevolution can arise when reciprocal evolutionary change happens between interacting parties, driving or impeding the diversification of the interacting clades (Yoder and Nuismer, 2010; Hembry et al., 2014). Coevolution can leave detectable signals in the cophylogenetic patterns of bipartitie species interactions, such as those formed by hosts and their mutualistic, commensal, or parasitic symbionts. Significant effort has been devoted to studying topological congruence between phylogenies of interacting organisms as an indicator of cospeciation or codiversification (e.g., Legendre et al. 2002; de Vienne et al. 2007; Hoyal Cuthill and Charleston 2012; Hayward et al. 2021; Perez-Lamarque and Morlon 2024; reviewed in Nieberding and Olivieri 2007; de Vienne et al. 2013; Martínez-Aquino 2016; Dismukes et al. 2022). Recent cophylogenetic research has forayed into linking cophylogenetic patterns with their underlying eco-evolutionary processes (Blasco-Costa et al., 2021). Another area of active research is cophylogenetic reconstruction, aiming to reconstruct the joint evolutionary history of interacting clades by finding a parsimonious series of events that give rise to the observed cophylogenetic patterns (e.g., Merkle et al. 2010; Baudet et al. 2015; Sinaimeri et al. 2023; reviewed in Charleston and Libeskind-Hadas 2014).

However, the study of the macroevolutionary rates of cophylogenetic systems remains in its infancy. Macroevolutionary rates can be defined, for single clades, as speciation and extinction rates that can be inferred from phylogenetic systems (Stadler, 2013; Harmon et al., 2014). Here, we define macroevolutionary rates of species interactions (i.e., of interacting clades) as speciation and extinction rates that can be inferred from cophylogenetic systems. Although much recent work has been focused on the advancement of phylogenetic comparative methods for estimating the speciation, extinction, or net diversification rates of single clades (Garamszegi, 2014; Revell and Harmon, 2022; Morlon et al., 2024), single-clade-based phylogenetic methods cannot be readily used for cophylogenetic systems. Specifically, although these single-clade-based methods allow diversification rates to be inferred separately for host and symbiont clades in a given cophylogenetic system, these rate estimates cannot provide a full picture of the prevalence of different types of speciation in the cophylogeny (Charleston and Perkins, 2006; Charleston and Libeskind-Hadas, 2014). Motivated by a desire to better understand parasitic symbionts (i.e., pathogens), Alcala et al. (2017) represents an important advance in estimating macroevolutionary rates of species interactions from cophylogenetic systems. This approach focuses on estimating the rate of cospeciation and the probability of host switching in the symbiont clade. However, this approach has two major disadvantages when the entire cophylogenetic system, as opposed to the symbiont clade alone, is of interest: (i) it does not allow host speciation and symbiont speciation to occur on the same timescale; (ii) it does not take into account host extinction, in which symbionts relying on the extinct host goes extinct as well. A promising, more recent approach can already tease apart the relative contributions of different macroevolutionary processes (cospeciation, host switching, pollinator speciation, and pollinator extinction) in generating cophylogenetic patterns (Satler et al., 2019). However, this method suffers the same problem of inconsistent timescales between host and symbiont diversification, resulting in zero rate estimates for host switching and cospeciation in some cases. Fortunately, recent advances have allowed simulating cophylogenetic systems as a result of constant-rate birth-death processes, simultaneously incorporating speciation and extinction in both the host and symbiont clades (Dismukes and Heath, 2021). This new model has paved the way for developing new statistical methods for estimating macroevolutionary rates of bipartite species interactions.

Approximate Bayesian Computation (ABC) has been shown to be a useful tool for estimating macroevolutionary rates of species interactions from cophylogenetic systems (Alcala et al., 2017). ABC is a class of simulation-based inference that has proven especially suitable for models whose likelihoods are impossible or difficult to obtain (Marin et al., 2012; Sunnåker et al., 2013). Initially developed by geneticists, ABC has been increasingly used in ecology and evolution (Pantel and Becks, 2023). In the most basic form of ABC, real data is compared to numerous simulations generated using different parameter samples, and parameters samples that generate simulations similar enough to the real data constitute the approximate posterior distribution of that parameter; the key to successful ABC is designing informative summary statistics that are informative and low in dimensionality (Sunnåker et al., 2013). Conceptual frameworks and standard practices have been established for using approximate Bayesian computation for both parameter estimation and model selection in ecology and evolution (Csilléry et al., 2010, 2012; Janzen et al., 2015; Pontarp et al., 2019; Pantel and Becks, 2023). As for estimating macroevolutionary rates of species interactions from cophylogenetic systems, the existing ABC approach shows promising potential but also suffers significant difficulty in implementation (Alcala et al., 2017). Specifically, the implementation of this approach requires highly customized representation of cophylogenies, a multitude of network summary statistics, dimension reduction techniques, and machine learning techniques. An easy-to-implement ABC approach to estimating macroevolutionary rates of species interactions from cophylogenies is yet to be developed.

Here, we develop a new ABC approach that, while taking into account host and symbiont extinction, infers the rates of four types of speciation processes from an extant cophylogeny: (i) host speciation, (ii) symbiont speciation without host-switching, (iii) symbiont speciation with host-switching, and (iv) cospeciation. This ABC approach is easy to implement thanks to our newly designed summary statistics, which take into account properties of the cophylogeny that are both related to or irrespective of the sizes of the host and symbiont trees (i.e., size-based and size-free summary statistics). We show that the performance of the parameter inference remains reasonably good despite the “curse of dimensionality”, process stochasticity of speciation and extinction, and partial information.

## 2 Methods

In brief, we follow these two steps to estimate the speciation rates of a target cophylogeny: (i) we simulate cophylogenies following *a birth-death model for co-diversification*. (ii) In an *ABC framework*, the simulated phylogenies are compared to the target cophylogeny to obtain estimates of the speciation rates. Notably, the comparison between the simulated and target cophylogenies is decided by *summary statistics*. In our *performance evaluation*, we perform this two-step ABC procedure, using cophylogenies simulated with known parameter values as the target cophylogeny. In our *application to empirical data*, we perform the same two-step ABC procedure using a cophylogeny compiled from empirical data as the target cophylogeny. Below, we provide details of each of these components.

### 2.1 A Birth-Death Model for Co-diversification

We use the sim_cophyBD function from the R package treeducken (ver. 1.1.0; Dismukes and Heath 2021) for simulating cophylogenies following a birth–death model. The model in treeducken allows the simulation of host and symbiont co-diversification. The model considers six types of speciation or extinction events (Fig. 1): host sepeciation (*λ*_*H*_), symbiont speciation without host switching (*λ*_*S*_), cospeciation (*λ*_*C*_), symbiont speciation with host switching(*λ*_*W*_), host extinction (*µ*_*H*_), and symbiont extinction (*µ*_*S*_). Each type of event is controlled by a rate parameter that represents the number of events per unit time following a Poisson process. For example, a rate of 1 means one event occurs per unit time in the cophylogenetic system.

**Figure 1:**
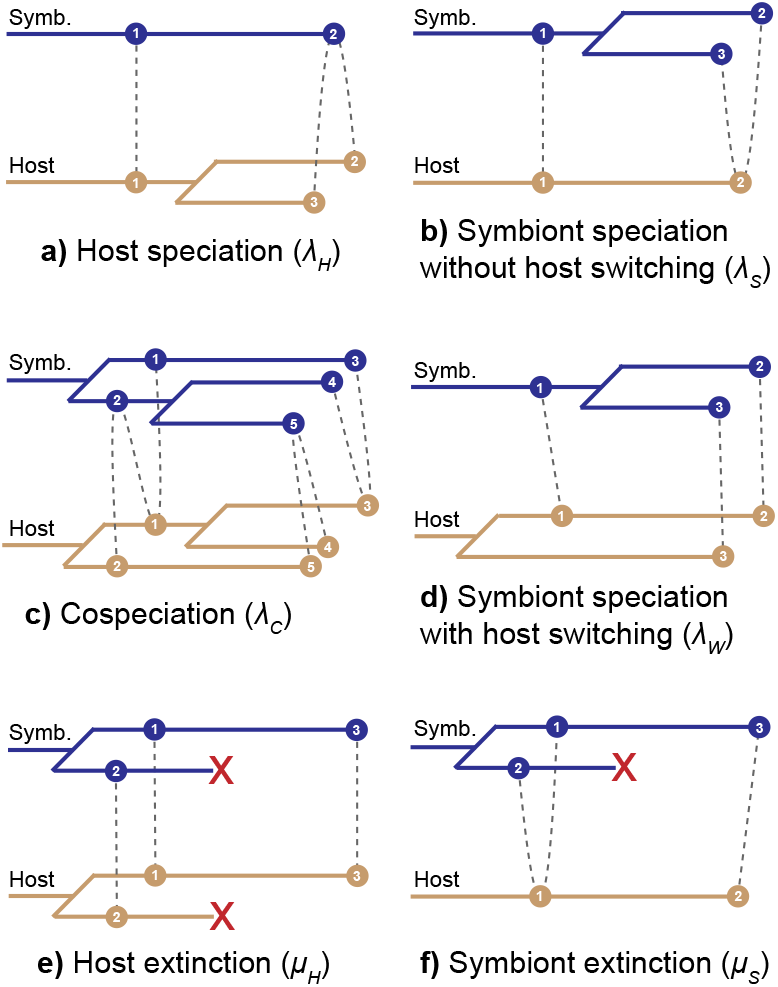
Speciation and extinction events giving rise to cophylogenetic patterns. Macroevolutionary rates of species interactions considered in this study (denoted by Greek letters in parentheses) are defined as, for each type of event, the number of events per unit time. **(a)** Host speciation (*λ*_*H*_): both descendent hosts retain association with the original symbiont. **(b)** Symbiont speciation without host switching (*λ*_*S*_): both descendent symbionts retain association with the original host. **(c)** Cospeciation (*λ*_*C*_): a host and one of its symbionts, randomly selected, undergoes speciation simultaneously (host 1 into hosts 3 and 4; symbiont 2 into symbionts 4 and 5). Each descendant host forms association with one of the descendant symbionts (host 3 with symbiont 4; host 4 with symbiont 5). The ancestral host’s and symbiont’s remaining associations are randomly sorted among the descendants. **(d)** Symbiont speciation with host switching (*λ*_*W*_): same as (b), but here one of the descendant symbionts switches to a new, randomly selected hostt. **(e)** Host extinction (*µ*_*H*_): note that the symbiont(s) relying on this host goes extinct as well. **(f)** Symbiont extinction (*µ*_*S*_). These event definitions are identical to those in the simulation tool, except that the rate of symbiont speciation with host switching (*λ*_*W*_) is denoted by *χ* in the original study (Dismukes and Heath, 2021).

Because this approach is designed to infer macroevolutionary rates of species interactions from extant cophylogenetic systems, the extinction rate must not exceed the speciation rate for the host and symbiont clades. We further parameterize the model by defining relative extinction rates *ϵ*_*H*_ (for the host clade) and *ϵ*_*S*_ (for the symbiont clade) as follows,

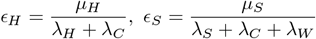

where both *ϵ*_*H*_ and *ϵ*_*S*_ range between 0 and 1. In the ABC framework (detailed below), we assume that *ϵ*_*H*_ and *ϵ*_*S*_ are known and aim to estimate the four speciation rates (*λ*_*H*_, *λ*_*S*_, *λ*_*C*_, *λ*_*W*_).

### 2.2 ABC Framework

Speciation rates (*λ*_*H*_, *λ*_*S*_, *λ*_*C*_, *λ*_*W*_) in a cophylogenetic system can be collectively seen as a multivariate parameter *θ*, which is treated as a random variable in Bayesian parameter inference. The posterior probability of *θ* given observed data *D* can be, in theory, obtained by

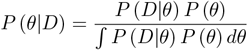

where *P* (*θ*) is known as the prior distribution and *P* (*D*| *θ*) as the likelihood function (i.e., the probability of observing the cophylogeny *D* given the parameter *θ*). However, when *D* is a cophylogeny and *θ* is the parameter in the birth-death model, the likelihood function is difficult to obtain, making Approximate Bayesian Computation (ABC) an especially useful approach for parameter estimation (Alcala et al., 2017). An ABC approach does not explicitly compute the likelihood but instead uses simulations to approximate the posterior distribution. Here, we use the basic form of a rejection-based ABC approach (Csilléry et al., 2012) following these steps. First, cophylogenies are simulated with the birth-death model (see the previous section in Methods) using samples of *θ* drawn from the prior distribution. Then, each sample from the prior distribution is accepted if the distance between the resulting simulated cophylogeny and *D* is below a given threshold. Alternatively, a sample is rejected if the distance is above the threshold. The distances are determined by summary statistics (see the following section in Methods). The threshold is decided by the tolerance rate, which determines the proportion of simulations with the smallest distance to accept. Eventually, accepted samples of *θ* constitute the approximate posterior distribution.

### 2.3 Summary Statistics

The key to successful ABC parameter inference is summary statistics that decide the distance between a simulation and the observed data (Sunnåker et al., 2013). A cophylogenetic system (e.g., Fig. 2a) consists of two phylogenies and the network formed by the tips of the two trees. In theory, one could use indices of phylogenies and networks as the summary statistics for ABC. Indices for phylogenies include phylogenetic diversity (Clarke and Warwick, 2001), the gamma statistic (Pybus and Harvey, 2000) and Sackin index measuring tree shape (Sackin, 1972). Indices for networks include global properties such as nestedness, modularity, connectance (Guimaraes Jr, 2020), and meso-scale properties such as motif frequencies (Simmons et al., 2019). Indices concerning both the phylogeny and network components of a cophylogeny include the mantel correlation, a measure of phylogenetic conservatism of bipartite species association (Maliet et al., 2020). However, single indices tend not to be informative enough as summary statistics for phylogenetic systems (Janzen et al., 2015; Janzen and Etienne, 2024). On the other hand, when we combine multiple of these indices in our preliminary exploratory analyses, parameter convergence is generally difficult. This is unsurprising because the high dimensionality of summary statistics can introduce the “curse of dimensionality”, that is, the match between a simulation and the observed data gets significantly more unlikely when the dimensionality of summary statistics increases (Sunnåker et al., 2013). The difficulty in convergence in our preliminary analyses also agrees with the previous study on estimating macroevolutionary rates of species interactions from cophylogenies, where accurate parameter inference based on numerous indices is difficult without dimension reduction and machine learning techniques (Alcala et al., 2017). Because of these reasons, we do not choose index-based summary statistics for our approach.

**Figure 2:**
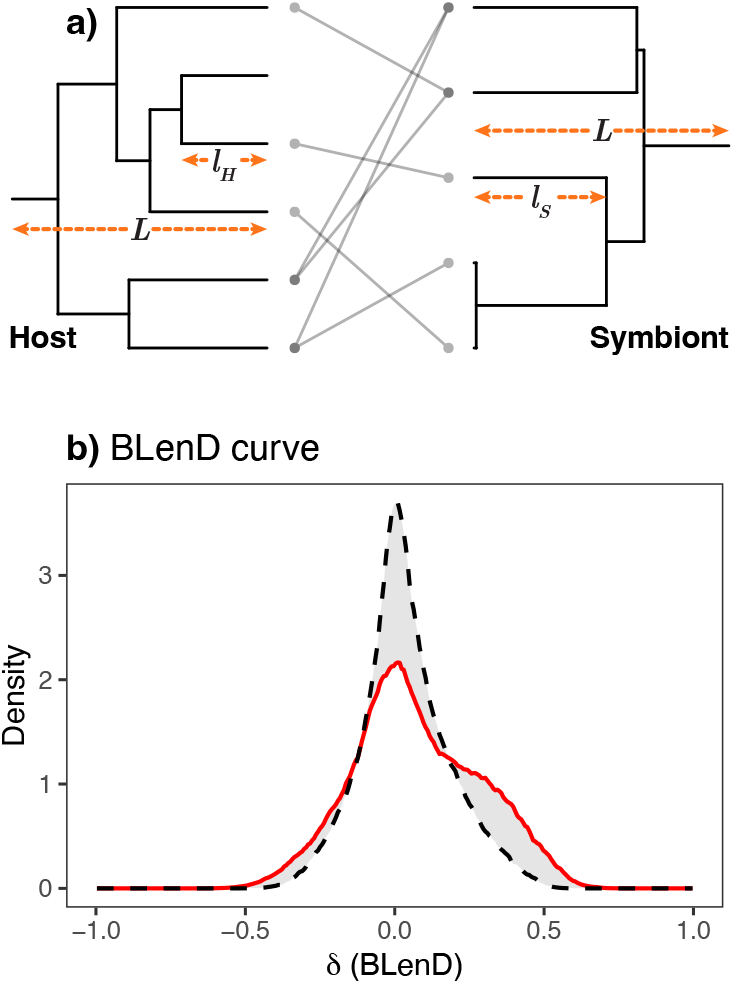
The Branch Length Difference (BLenD) curve as summary statistics for approximate Bayesian computation. **(a)** For each association (line with dotted ends) in the cophylogeny, we measure three lengths: the branch length of the host tip *l*_*H*_, the branch length of the symbiont tip *l*_*S*_, and the total length of the cophylogeny *L* (i.e., assumed to the same for the host and symbiont phylogenies). Then, we obtain *δ* = (*l*_*H*_ − *l*_*S*_)*/L* for every association in the cophylogeny. This allows us to estimate the density of *δ* in the entire cophylogeny (i.e., the BLenD curve) **(b)** Cophylogenies A (solid line) and B (dashed line) with different speciation rates have different BLenD curves. In both Cophylogenies A and B, *ϵ*_*H*_ = *ϵ*_*S*_ = 0.3. In Cophylogeny A, *λ*_*H*_ = 0.9, *λ*_*S*_ = 0.3, *λ*_*C*_ = 0.4, *λ*_*W*_ = 1.7; in Cophylogeny B, *λ*_*H*_ = 1.1, *λ*_*S*_ = 1.2, *λ*_*C*_ = 0.7, *λ*_*W*_ = 0.3. For each curve, the number of replicates is 500 and the median density across all replicates is used.

As opposed to index-based summary statistics, normalized curves have been proposed as an alternative type of summary statistics for ABC. Specifically, a pioneering study has used a normalized lineage-through-time curve (nLTT), which is irrespective of the size or length of the phylogenetic tree, to effectively estimate constant speciation rates from single phylogenies while taking into account constant-rate lineage extinction (Janzen et al., 2015). The nLTT summary statistics are informative, computationally efficient, and not obviously affected by the “curse of dimensionality”. Inspired by this work, here we adapt the use of normalized curves as summary statistics for cophylogenetic systems, the details of which are given below.

#### 2.3.1 BLenD Curve

The BLenD (Branch Length Difference) curve is a normalized curve irrespective of the sizes of the cophylogeny of interest. For any association (between a host and a symbiont) in the cophylogeny, it is possible to obtain

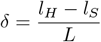

where *l*_*H*_ is the branch length of the host tip, and *l*_*S*_ is the branch length of the symbiont tip, and *L* is the length (or height) of the cophylogeny from its root to tips (Fig. 2). We refer to *δ* (− 1 *< δ <* 1) as the normalized Branch Length Difference (BLenD). 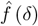,the density function of *δ*, can then be estimated from all observed values of *δ* in the cophylogeny (*δ*_1_, *δ*_2_, …, *δ*_*n*_, where *n* is the total number of associations; each association is shown as a line with dotted ends in Fig. 2). This estimation is implemented as kernel density estimation using the function `density(kernel = “gaussian”)` in the `stats` package in R (ver. 4.4.0; R Core Team 2024). We hereafter refer to 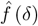 as the BLenD curve.

Now, let us imagine that we have two cophylogenies instead of one, which we will call Cophylogeny A and Cophylogeny B. Thus, the distance between Cophylogeny A and Cophylogeny B can be defined as:

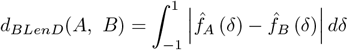

where 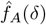 and 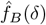 are the BLenD curves for Cophylogenies A and B, respectively. *d*_*BLenD*_(*A, B*) is equal to area between the BLenD curves of Cophylogenies A and B (shaded area in Fig. 2). Similarly to Janzen et al. (2015), this definition of distance achieves a natural weighing of the contribution of all the 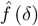-differences between Cophylogenies A and B; also similarly, the BLenD curve is implemented as a vector of summary statistics. In effect, *d*_*BLenD*_(*A, B*) is equal to zero only when the two BLenD curves compared are identical.

#### 2.3.2 Tree Sizes

In addition to the newly designed the BLenD curve, we consider the sizes of the two trees (the numbers of tips in the host and symbiont trees) in the cophylogeny as summary statistics because speciation and extinction rates will almost certainly affect the numbers of tips in the host and symbiont phylogenies. The distance between Cophylogenies A and B, based on the tree sizes of their *host* phylogenies, can be defined as:

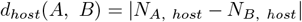

where *N*_*A*, *host*_ and *N*_*B*, *host*_ are the sizes of the host phylogenies in Cophylogenies A and B, respectively. Similarly, the distance between Cophylogenies A and B, based on the tree sizes of their *symbiont* phylogenies, can be defined as:

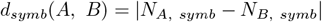

where *N*_*A*, *symb*_ and *N*_*B*, *symb*_ are the sizes of the symbiont phylogenies in Cophylogenies A and B, respectively. Tree sizes are log-transformed before being used to calculate tree-size-based distances.

#### 2.3.3 Combining Summary Statistics

Combining different summary statistics in general increases the informativeness of summary statistics for ABC (Sunnåker et al., 2013), so here we test whether BLenD curve and tree sizes combined can achieve better inference results than either of them alone. When the BLenD curve and tree sizes are used in tandem, we standardize the summary statistics by dividing them by their standard deviation to ensure that the BLenD curve and each of the two tree size statistics have comparable variation among simulations. The distance between Cophylogenies A and B is computed as the sum of the BLenD-based and tree-size-based distances.

### 2.4 Performance Evaluation

We are particularly interested in testing this ABC approach’s performance with cophylogenies with rate heterogeneity between the four types of speciation (*λ*_*H*_, *λ*_*S*_, *λ*_*C*_, *λ*_*W*_). We focus our testing on the most basic form of rate heterogeneity, that is, one type of speciation occurs at a higher rate than the other three types. This most basic form of rate heterogeneity includes four possibilities: *λ*_*H*_ *>λ*_*S*_=*λ*_*C*_=*λ*_*W*_, *λ*_*S*_*>λ*_*H*_ =*λ*_*C*_=*λ*_*W*_, *λ*_*C*_*>λ*_*H*_ =*λ*_*S*_=*λ*_*W*_, or *λ*_*W*_ *>λ*_*H*_ =*λ*_*S*_=*λ*_*C*_. They correspond to situations where only one type of speciation predominates in the cophylogenetic system. To test the accuracy of the parameter estimator, we perform convergence analyses to test whether the approximate posterior distribution approaches the true parameter values when tolerance rate approaches zero. We document both the parameter estimates and their residuals (i.e., estimates minus the true value).

Besides the parameter inference problem, we also evaluate how correctly our ABC approach can *categorically* detect a specific type of speciation that occurs at a higher rate than the other three types. A correct detection is considered to have been achieved when the type of speciation with the highest true rate also has the highest median rate estimate in the approximate posterior distribution. We evaluate how the detection correctness of this ABC approach compares to a random guess. To generate simulated cophylogenies for five combinations of *ϵ*_*H*_ and *ϵ*_*S*_ (0/0, 0.3/0.3, 0.7/0, 0/0.7, 0.7/0.7 for *ϵ*_*H*_ */ ϵ*_*S*_), we run 50000 simulations each containing 100 replicates and take the median BLenD density and tree sizes across all replicates.

### 2.5 Application to Empirical Data

The ABC approach is applicable to cophylogenetic datasets where the host and symbiont phylogenies are both time-calibrated and have the same stem age. This requirement is important because the timescale has to be held consistent for all six of the speciation/extinction processes (Fig. 1). One empirical dataset that meets this requirement is a dataset of Batesian mimicry compiled by Van Dam et al. (2024), where beetles in the genus *Doliops* mimic beetles in the genus *Pachyrhynchus*, both native to the Philippines. This mimicry can be seen as a case of commensalism, which is defined as a type of interaction where one party in the interaction receives a benefit while the other party receives neither a cost nor a benefit (Bronstein, 2015). Particularly in this case, the mimics (*Doliops*) receive a protective benefit while the models (*Pachyrhynchus*) are largely unaffected. Host-symbiont models have been fitted to this dataset in the original study (Van Dam et al., 2024), with mimics considered as symbionts and models as hosts. The stem age for both *Pachyrhynchus* and *Doliops* has been geologically calibrated for the maximum age of the Philippines and estimated to be 25-30 Myr in the original study. We use 27.5 Myr, the midpoint of this range, as the total length of both phylogenies (distance from the root to tips). We used the `phylo.tracer` function from the `physketch` package (Revell, 2020) to extract the two beetle phylogenies from Figure 3 of the original study. We then code the association between species manually based on Figure 2 of the original study. For the beetle cophylogeny to be comparable with cophylogenies simulated with treeducken, we rescale the beetle cophylogeny such that the total length of the beetle cophylogeny is equal to the total length of the simulated cophylogenies (i.e., 27.5 Myr on the realistic timescale is convertible to 2 unit time on the `treeducken` timescale as defined in Dismukes and Heath 2021).

**Figure 3:**
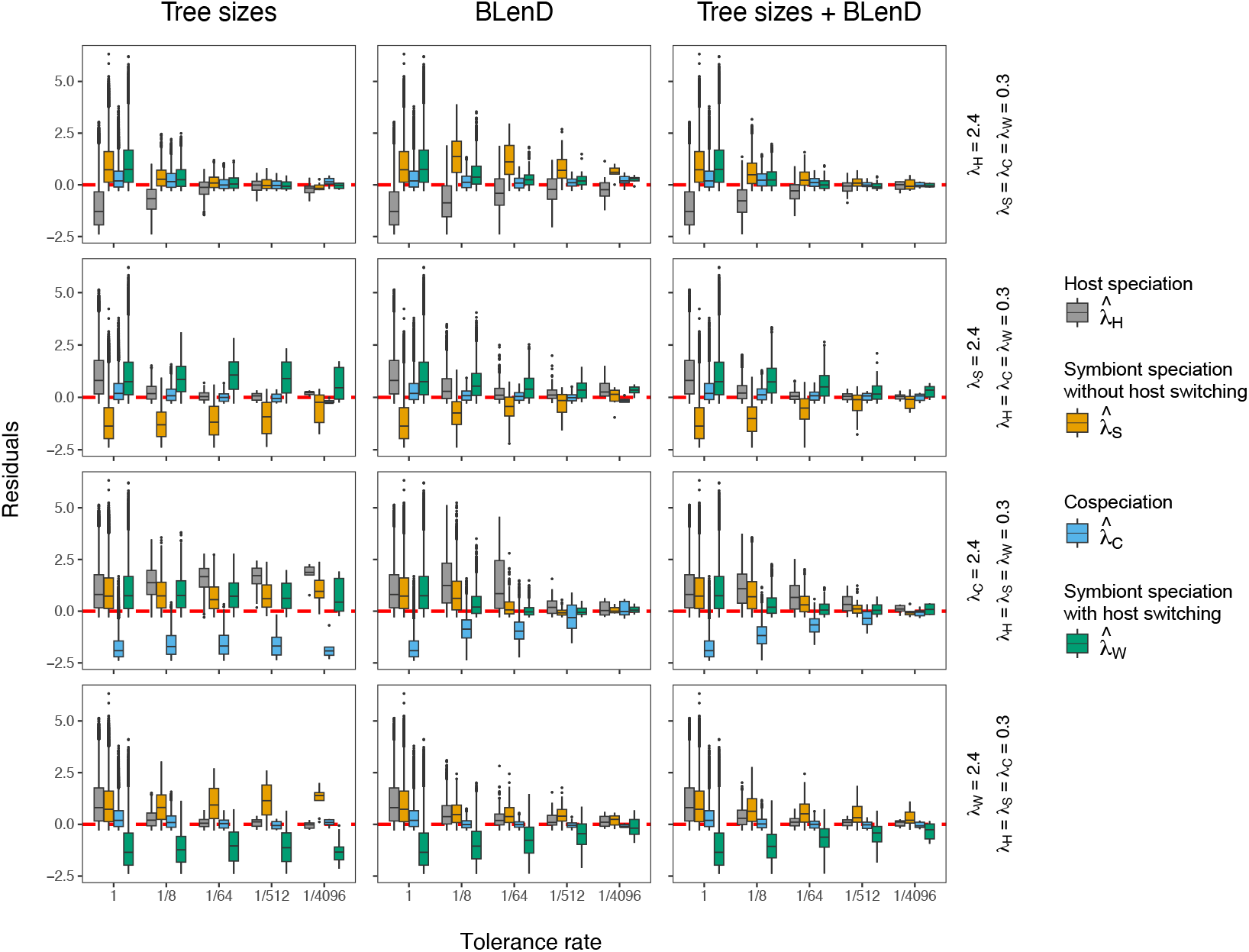
Convergence plots of the speciation rate estimates 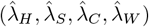 when *ϵ*_*H*_ = *ϵ*_*S*_ = 0.3. Residuals are calculated as estimates minus the true values (*λ*_*H*_, *λ*_*S*_, *λ*_*C*_, *λ*_*W*_). The estimates converge well if the residuals approach zero as tolerance rate approaches zero. Simulated cophylogenies whose true parameters are known (vertical text on the right) are used as observed data in the ABC framework. Each observed cophylogeny contains 500 replicates; the median BLenD density and tree sizes are taken across all replicates before being used as observations in the ABC. Convergence plots under other assumptions of *ϵ*_*H*_ and *ϵ*_*S*_ are presented in Figs. S1-S4.

For this empirical cophylogenetic dataset, we estimate the four speciation rates (*λ*_*H*_, *λ*_*S*_, *λ*_*C*_, *λ*_*W*_) using the same ABC framework (see a previous section in Methods) and the same sets of simulated cophylogenies that we use for performance evaluation (see a section in Methods). As a necessary step when applying ABC to real data (Pantel and Becks, 2023), we also perform posterior predictive checks of model fit to see how well the simulatied cophylogenies that are accepted in the ABC framework (i.e., those that contribute to the approximate posterior distribution of the parameter estimates) resemble the beetle cophylogeny.

## 3 Results

### 3.1 Performance

#### 3.1.1 Rate Estimation

Here we focus on how different choices of summary statistics compare in terms of convergence. Good convergence is considered to have been achieved when both the precision (i.e., how narrow the approximate posterior distribution is) and accuracy (i.e., how close the approximate posterior distribution is to the true value) of parameter estimation are high. Our convergence analyses reveal that, across all combinations of relative extinction rate *ϵ*_*H*_ and *ϵ*_*S*_ that we consider, parameter estimates generally converge to the true values of the four lambdas (*λ*_*H*_, *λ*_*S*_, *λ*_*C*_, *λ*_*W*_) best when a combination of tree sizes and the BLenD curve are used as summary statistics (the third column compared to the first and second columns in Figs. 3, S1-S4). When only tree sizes are used (the first column in Figs. 3, S1-S4), the parameter estimates converge well only when *λ*_*H*_ is the highest among the four lambdas; otherwise the parameter estimates converge poorly, having a low precision or low accuracy. When only the BLenD curve is used (the second column in Figs. 3, S1-S4), the parameter estimates converge well when *λ*_*S*_, *λ*_*C*_, or *λ*_*W*_ is the highest among the four lambdas; however, the parameter estimates converge poorly when *λ*_*H*_ is the highest among the four lambdas. Thus, in terms of convergence, tree sizes alone perform well when the BLenD curve alone performs poorly; conversely, tree sizes alone perform poorly when the BLenD curve alone performs well. The performance of a combination of tree sizes and the BLenD curve (the third column in Figs. 3, S1-S4) is generally on par with the better of the two (tree sizes alone and the BLenD curve alone) and, in some cases, outperforms the better of the two (Figs. S1 & S2).

The combination of tree sizes and the BLenD curve is also shown to be generally the best at handling different types of rate heterogeneity and the least sensitive to relative extinction rates (Table 1, S1 & S2). The tree sizes, when used alone, are especially poor at handling rate heterogeneity where *λ*_*C*_ is the highest among the four lambdas (Table S2), but this type of rate heterogeneity is handled well by a combination of tree sizes and the BLenD curve (Table 1). Although the BLenD curve alone performs relatively well with most types of rate heterogeneity (Table S2), it is outperformed by a combination of tree sizes and the BLenD curve when *λ*_*H*_ is the highest among the four lambdas and the *ϵ*_*S*_ is as high as 0.7 (Table 1).

**Table 1:** Performance of speciation rate estimation and speciation rate heterogeneity detection under different assumptions of relative extinctions rates (*ϵ*_*H*_ and *ϵ*_*S*_). Four rate parameters are considered: *λ*_*H*_ - host speciation; *λ*_*S*_ - host speciation without host switching; *λ*_*C*_ - cospeciation; *λ*_*W*_ - symbiont speciation with host switching.

| Relative extinction rates |  | True speciation rates<sup>a</sup> |  |  |  | Rate estimates<sup>b</sup> |  |  |  | Rate heterogeneity detection |  |
| --- | --- | --- | --- | --- | --- | --- | --- | --- | --- | --- | --- |
| $\epsilon_H$ | $\epsilon_S$ | $\lambda_H$ | $\lambda_S$ | $\lambda_C$ | $\lambda_W$ | $\hat{\lambda}_H$ | $\hat{\lambda}_S$ | $\hat{\lambda}_C$ | $\hat{\lambda}_W$ | Correctness<sup>c</sup> | Improvement<sup>d</sup> |
| 0 | 0 | **2.4** | 0.3 | 0.3 | 0.3 | **2.19 (0.12)** | 0.41 (0.18) | 0.28 (0.12) | 0.14 (0.1) | 96% | 3.832 |
| 0 | 0 | 0.3 | **2.4** | 0.3 | 0.3 | 0.26 (0.16) | **2.3 (0.16)** | 0.3 (0.15) | 0.21 (0.14) | 72% | 2.88 |
| 0 | 0 | 0.3 | 0.3 | **2.4** | 0.3 | 0.42 (0.19) | 0.22 (0.12) | **2.17 (0.15)** | 0.12 (0.09) | 94% | 3.752 |
| 0 | 0 | 0.3 | 0.3 | 0.3 | **2.4** | 0.22 (0.12) | 0.75 (0.77) | 0.34 (0.18) | **1.74 (0.67)** | 62% | 2.48 |
| 0.3 | 0.3 | **2.4** | 0.3 | 0.3 | 0.3 | **2.35 (0.25)** | 0.33 (0.3) | 0.3 (0.12) | 0.28 (0.09) | 96% | 3.856 |
| 0.3 | 0.3 | 0.3 | **2.4** | 0.3 | 0.3 | 0.33 (0.17) | **2.2 (0.4)** | 0.3 (0.18) | 0.58 (0.31) | 47% | 1.896 |
| 0.3 | 0.3 | 0.3 | 0.3 | **2.4** | 0.3 | 0.41 (0.2) | 0.24 (0.2) | **2.32 (0.19)** | 0.39 (0.25) | 70% | 2.808 |
| 0.3 | 0.3 | 0.3 | 0.3 | 0.3 | **2.4** | 0.35 (0.16) | 0.64 (0.48) | 0.27 (0.15) | **2.05 (0.44)** | 68% | 2.736 |
| 0.7 | 0 | **2.4** | 0.3 | 0.3 | 0.3 | **2.45 (0.2)** | 0.4 (0.24) | 0.24 (0.16) | 0.32 (0.22) | 96% | 3.856 |
| 0.7 | 0 | 0.3 | **2.4** | 0.3 | 0.3 | 0.3 (0.11) | **2.1 (0.27)** | 0.28 (0.14) | 0.59 (0.3) | 42% | 1.696 |
| 0.7 | 0 | 0.3 | 0.3 | **2.4** | 0.3 | 0.47 (0.26) | 0.53 (0.4) | **2.16 (0.29)** | 0.27 (0.12) | 64% | 2.568 |
| 0.7 | 0 | 0.3 | 0.3 | 0.3 | **2.4** | 0.29 (0.12) | 0.79 (0.68) | 0.35 (0.12) | **1.85 (0.73)** | 65% | 2.592 |
| 0 | 0.7 | **2.4** | 0.3 | 0.3 | 0.3 | **2.45 (0.14)** | 0.41 (0.15) | 0.28 (0.12) | 0.28 (0.12) | 98% | 3.936 |
| 0 | 0.7 | 0.3 | **2.4** | 0.3 | 0.3 | 0.26 (0.16) | **1.96 (0.44)** | 0.34 (0.12) | 0.65 (0.39) | 51% | 2.056 |
| 0 | 0.7 | 0.3 | 0.3 | **2.4** | 0.3 | 0.33 (0.44) | 0.35 (0.17) | **2.37 (0.36)** | 0.36 (0.25) | 69% | 2.776 |
| 0 | 0.7 | 0.3 | 0.3 | 0.3 | **2.4** | 0.44 (0.18) | 0.78 (0.52) | 0.22 (0.17) | **1.85 (0.47)** | 66% | 2.656 |
| 0.7 | 0.7 | **2.4** | 0.3 | 0.3 | 0.3 | **2.36 (0.19)** | 0.42 (0.26) | 0.3 (0.12) | 0.23 (0.16) | 98% | 3.904 |
| 0.7 | 0.7 | 0.3 | **2.4** | 0.3 | 0.3 | 0.26 (0.19) | **1.71 (0.74)** | 0.41 (0.21) | 0.86 (0.57) | 45% | 1.792 |
| 0.7 | 0.7 | 0.3 | 0.3 | **2.4** | 0.3 | 0.63 (0.26) | 0.42 (0.25) | **2.14 (0.22)** | 0.34 (0.21) | 58% | 2.336 |
| 0.7 | 0.7 | 0.3 | 0.3 | 0.3 | **2.4** | 0.25 (0.16) | 0.77 (0.72) | 0.34 (0.17) | **1.72 (0.65)** | 65% | 2.608 |

Note: tree sizes and the BLenD curve are used in tandem as summary statistics. The tolerance rate used is 1/4096. See Tables S1 & S2 for results for when tree sizes or the BLenD curve are used alone.
<sup>a</sup> Fixed, known parameter values used to generate the observed data (cophylogeny) in the ABC framework. 500 replicates are generated for each set of parameter values.
<sup>b</sup> Means (outside parentheses) and standard deviations (inside parentheses) of the rate estimates. The median BLenD density and tree sizes are taken across all 500 replicates of the observed data before being used in the ABC.
<sup>c</sup> Percentage of the 500 replicates of observed data for which the highest rate among $\lambda_H, \lambda_S, \lambda_C, \lambda_W$ is detected correctly.
<sup>d</sup> Actual correctness (see footnote c) divided by the expected correctness of a random guess among $\lambda_H, \lambda_S, \lambda_C, \lambda_W$ (25%).

#### 3.1.2 Rate Heterogeneity Detection

As for the detection of speciation rate heterogeneity, combining BLenD and tree sizes as summary statistics, again, tends to outperform the use of the BLenD curve or tree sizes alone (Table 1, S1 & S2). We are specifically interested in identifying the highest rate among the four lambdas *λ*_*H*_, *λ*_*S*_, *λ*_*C*_, *λ*_*W*_, and a random guess among the four lambdas is expected to have a 25% correctness. An improvement value (i.e., the actual correctness divided by 25%) greater than one suggests that the inference performs better than a random guess, and vice versa. Our results show that the tree sizes alone perform consistently worse than the random guess when *λ*_*C*_ is the highest among the four lambdas (Table S1), rendering the tree sizes unsuitable summary statistics for this purpose. The use of the BLenD curve alone performs better than a random guess in most cases (Table S2), but the performance is much improved when the BLenD curve is used in tandem with tree sizes (Table 1).

When the tree sizes and BLenD curve are used in tandem, the detection of some types of rate heterogeneity is less sensitive to extinction than others (Table 1). The detection correctness, when *λ*_*H*_ is the highest among the four lambdas, remains high (96-98%) regardless of *ϵ*_*H*_ and *ϵ*_*S*_. Similarly, the detection correctness when *λ*_*W*_ is the highest among the four lambdas remains somewhat high (62-68%) regardless of *ϵ*_*H*_ and *ϵ*_*S*_. However, the detection correctness for the other two types of rate heterogeneity is more sensitive to *ϵ*_*H*_ and *ϵ*_*S*_. Notably, the detection correctness of rate heterogeneity where *λ*_*S*_ or *λ*_*C*_ is the highest among the four lambdas drops by 24-25% (from 72% to 47% and from 94% to 70%, respectively) when *ϵ*_*H*_ and *ϵ*_*S*_ increase from 0 to 0.3 (Table 1).

### 3.2 Rates Inferred from Beetle Mimicry

Across all the combinations of *ϵ*_*H*_ and *ϵ*_*S*_ that we consider, the ABC infers host speciation (*λ*_*H*_) as the type of speciation with the highest rate (Figs. 4 & S5; Table 2). We infer the mean rate of host speciation to be 1.80-3.22 events per unit time, while the the rates of symbiont speciation (with or without host switching) and cospeciation are inferred to be below 1 event per unit time. In terms of mean rate estimates, host speciation is 4.3-5.1 times faster than the median among all four types of speciation. Regardless of assumptions of *ϵ*_*H*_ and *ϵ*_*S*_, host speciation is consistently inferred to have the highest rate among the four types of speciation. Our performance evaluation has shown that the ABC approach generally performs well with rate heterogeneity where the host speciation rate is much higher than the other three types of speciation rates (see the Performance section in Results). In light of these performance evaluation results, the rate estimates from the beetle mimicry dataset suggest that host speciation is the fastest among all four types of speciation (i.e., host specieation, symbiont speciation without host switching, cospeciation, and symbiont speciation with host switching) in the cophylogenetic system. Our posterior predictive checks how that model fit remains reasonably good regardless of assumptions of relative extinction rates *ϵ*_*H*_ and *ϵ*_*S*_ (Figs. 4 & S5).

**Table 2:** Rate estimates 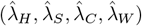 from the beetle mimicry dataset under different assumptions of *ϵ*_*H*_ and *ϵ*_*S*_.

| $\epsilon_H$ | $\epsilon_S$ | $\hat{\lambda}_H$ | $\hat{\lambda}_S$ | $\hat{\lambda}_C$ | $\hat{\lambda}_W$ |
| --- | --- | --- | --- | --- | --- |
| 0 | 0 | 1.80 (0.21) | 0.36 (0.21) | 0.48 (0.12) | 0.25 (0.19) |
| 0.3 | 0.3 | 2.42 (0.27) | 0.65 (0.43) | 0.42 (0.18) | 0.30 (0.22) |
| 0.7 | 0 | 3.22 (0.33) | 0.98 (0.43) | 0.34 (0.18) | 0.35 (0.28) |
| 0 | 0.7 | 2.12 (0.26) | 0.38 (0.27) | 0.46 (0.16) | 0.43 (0.22) |
| 0.7 | 0.7 | 3.21 (0.42) | 0.81 (0.52) | 0.35 (0.26) | 0.45 (0.27) |

Note: for the lambdas, means are outside and standard deviations are inside parentheses. For comparability with Table 1, all lambdas are *not* converted to events/Myr.

**Figure 4:**
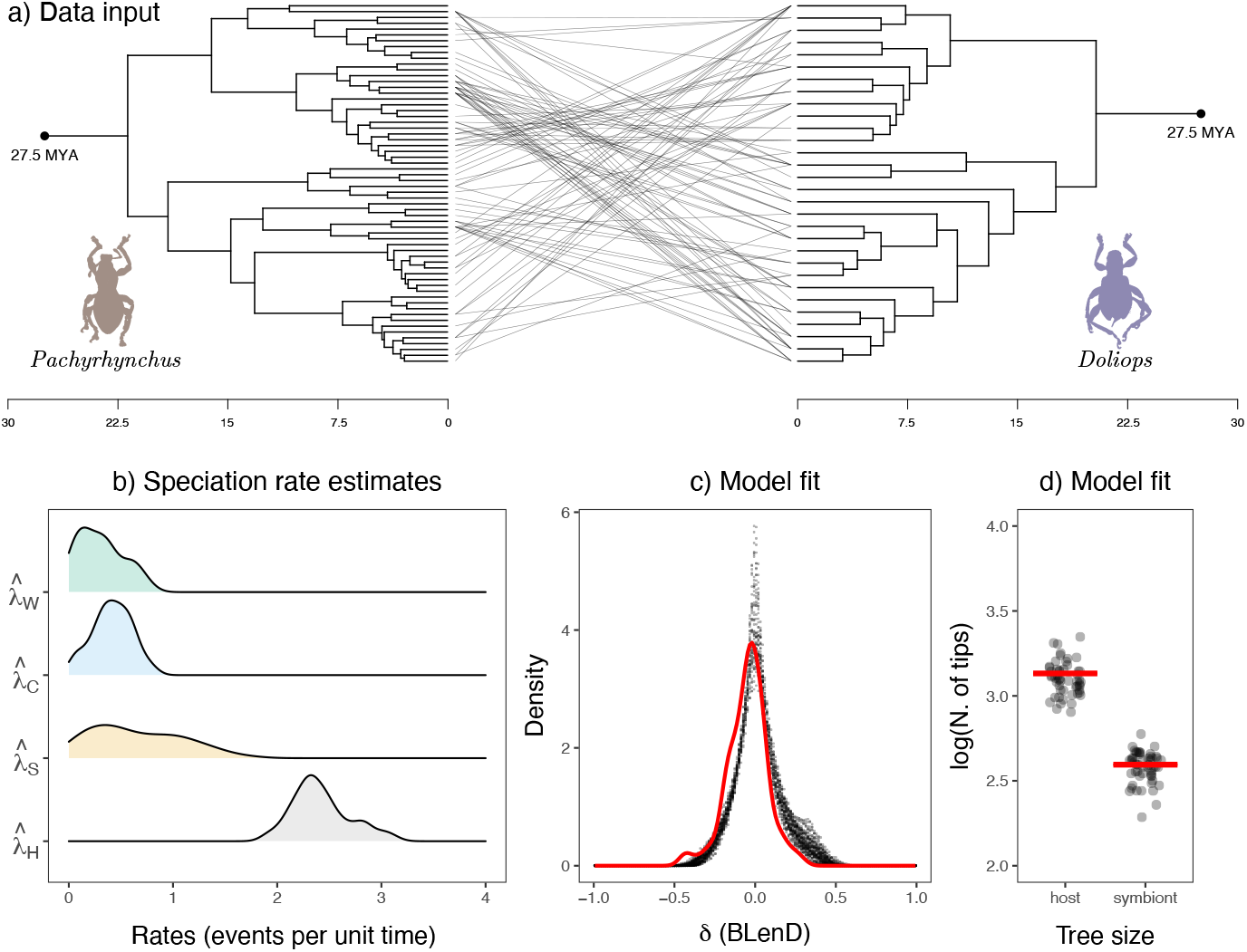
Speciation rate inference from the cophylogenetic dataset of beetle mimicry (Van Dam et al. 2024). Four rate parameters are considered: *λ*_*H*_ - host speciation; *λ*_*S*_ - host speciation without host switching; *λ*_*C*_ - cospeciation; *λ*_*W*_ - symbiont speciation with host switching. The BLenD curve and tree sizes are used in tandem as summary statistics. *ϵ*_*H*_ and *ϵ*_*S*_ are assumed to be 0.3. The tolerance rate used is 1/512. **(a)** Time-calibrated cophylogeny of *Pachyrhynchus* (models, treated as hosts) and *Doliops* (mimics, treated as symbionts). **(b)** Density curves of the speciation rate estimates. **(c & d)** Posterior predictive checks of model fit. Shown here are the BLenD curve and tree sizes of the beetle data (solid line) and accepted simulations in the ABC (dots). Results under other assumptions of *ϵ*_*H*_ and *ϵ*_*S*_ are presented in Fig. S5 and Table 2.

## 4 Discussion

Here we present a proof of concept that an ABC approach can be useful for inferring, simultaneously, rates of different types of speciation from a cophylogenetic system. Key to our approach is the newly designed summary statistics combining both size-based (tree sizes) and size-free statistics (the BLenD curve). The new approach presented here can be used to infer the predominant mode of speciation in a cophylogenetic system, which should be an important question to biologists interested in species interactions.

We show that the new ABC approach can handle several expected technical difficulties reasonably well. Specifically, (i) the “Curse of dimensionality” (Sunnåker et al., 2013): the ABC simultaneously infers the rates of four processes (four parameters). (ii) Stochasticity: both speciation and extinction are simulated as stochastic processes in the birth-death co-diversification model, introducing stochasticity to the resulting cophylogenies (Dismukes and Heath, 2021). (iii) Partial information: a cophylogeny contains only partial information because extinct lineages and historical association between hosts and symbionts are not known from extant cophylogenies. Compared to previous approachs to estimating macroevolutionary rates of species interactions from cophylogenies (Alcala et al., 2017; Satler et al., 2019), our new ABC approach allows speciation rate estimation under a more biologically realistic assumption that speciation-extinction dynamics occur simultaneously in the host and symbiont clades on the same timescale. In contrast to Alcala et al. (2017), the new approach achieves satisfactory accuracy without the use of highly customized representation of cophylogenies, dimension reduction, or machine learning.

### 4.1 Theoretical and Practical Considerations

Recent advances have revealed that extant phylogenetic systems are consistent with a myriad of speciation and extinction configurations (Louca and Pennell, 2020). Meaningful speciation rate inference would generally require *a priori* hypotheses about the extinction rates of the system of interest (Morlon et al., 2022, 2024). This identifiability issue likely can be extrapolated to cophylogenetic systems as well - in our beetle mimicry example, we find that model fit is not obviously better or worse under any particular assumption of relative extinction rates *ϵ*_*H*_ and *ϵ*_*S*_ (Figs. 4 & S5). However, rate estimates differ considerably under different assumptions of relative extinction rates *ϵ*_*H*_ and *ϵ*_*S*_ (Table 2). Thus, biologically realistic assumptions about the relative extinction rates of hosts and symbionts are critically important for speciation rate inference. Ideally, literature should be available on what might be a reasonable assumption of relative extinction rates. Such literature is likely system-specific (e.g., on figs and fig wasps, or specialized parasitisms) and requires expert knowledge about the system of interest. If such information is unavailable, the user should perform the same speciation rate inference under different assumptions of relative extincton rates to test the robustness of their conclusions.

We show that, as with phylogenies of single clades (Morlon et al., 2022, 2024), cophylogenies can still provide useful insights into diversification as long as realistic assumptions about extinction are made.

Given the numerous possibilities of rate heterogeneity between four types of speciation, we choose to focus on the most basic form of rate heterogeneity, one where one type of speciation predominates in the entire system. Testing the ABC approach on simulated datasets with diverse types of rate heterogeneity may extend the use of this approach. Other forms of rate heterogeneity may include, for example, situations where two (or three) types of speciation predominate. Those more complex forms of rate heterogeneity may not be dealt with as easily as those considered in this study; they would deserve to be the focus of a separate study. However, the most basic form of rate heterogeneity that we consider in this study may already be able to account for many observed patterns in nature, as we show with the beetle mimicry example (Fig. 4).

When the user performs this inference for an empirical system, it is usually not possible to know what the true parameter values are for their system of interest. Caution should be taken with the interpretation of results where the four lambda estimates 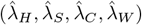 differ only slightly. Our performance test results show that, while large disparity between the four lambda estimates most likely reflects differences between their true rates (*λ*_*H*_, *λ*_*S*_, *λ*_*C*_, *λ*_*W*_), slight differences between the four lambda estimates can arise from errors of the ABC (Tables 1, S1 & S2). Thus, although strong disparity between the rate estimates of the four types of speciation likely reflects a biological reality, such conclusions may not be appropriate if the rate estimate differences are subtle.

In order to go beyond the proof of concept that we show here, technical improvements would be potentially useful. Single-layered feed-forward neural networks, a machine learning technique, have been used to improve the accuracy of ABC parameter inference (Csilléry et al., 2012). The improved accuracy is achieved through neural-network-based regression to correct rate estimates (Blum and François, 2010). This approach currently only works with a few single summary statistics. Adapting this regression-based approach for the use of normalized curves (such as BLenD) as summary statistics may be an interesting future direction. Another promising direction may be to enhance the ABC in this study with Sequential Monte Carlo (SMC), an algorithm that improves the accuracy of inference by iteratively generating new parameter samples from the posterior distribution and repeating the ABC procedure (for examples of ABC-SMC for (co)phylogenetic systems, see Janzen et al. 2015; Baudet et al. 2015; Sinaimeri et al. 2023).

As with previous cophylogenetic studies, our ABC cophylogenetic approach is not immune to errors due to phylogenetic reconstruction methods, incomplete taxon sampling, or discord between species trees and gene trees (e.g., Hughes et al. 2007; Onuferko et al. 2019). Additionally, apparent losses of symbionts in some host lineages may be the results of incomplete sampling of host-symbiont associations (Jackson and Charleston 2004; Charleston and Perkins 2006). For cophylogenetic studies in general, a statistical framework for quantifying these sources of error remains undeveloped but should be a rewarding future direction.

A related body of research has focused on reconciling the phylogeny of symbionts with that of their hosts (e.g., Merkle et al. 2010; Baudet et al. 2015; Sinaimeri et al. 2023; see Charleston and Perkins 2006; Charleston and Libeskind-Hadas 2014 for an overview). These studies map the symbiont phylogeny onto the host phylogeny to answer where cospeciations, duplications, host switches, and symbiont losses occur along branches of the host phylogeny. As an intermediate step toward such reconciliation, recent approaches have used ABC to estimate the frequencies of the aforementioned events (Baudet et al., 2015; Sinaimeri et al., 2023). It appears that one could, in theory, derive the rates of these events by dividing their frequencies by time. However, such derivation is impossible because of the lack of a timescale in these methods. Specifically, temporal constraints imposed by tree topologies (e.g., host switches are only possible between temporally coexisting species) are enforced in these methods only to the point where the *order* of events are feasible; this criterion is known as temporal feasibility (Stolzer et al., 2012). This means that these methods, in contrast to our approach, do not take into account branch lengths and are largely agnostic to the exact intervals between events, making it impossible to derive the rates of these events from their frequencies. Therefore, to the best of our knowledge, our study represents the first ABC cophylogenetic approach to offer a temporally explicit view of multiple types of macroevolutionary events.

### 4.2 Broader Implications

The user might want to interpret the results from our approach in light of results from other methods. For instance, the rate estimates from our approach offer valuable information about conservatism, a popular topic in the biology of species interactions (Gómez et al., 2010). In symbiont speciation without host switching (*λ*_*S*_), the descendant symbiont lineages retain association with the host itself. Differently, in cospeciation (*λ*_*C*_), the descendant symbiont lineages retain association with the descendants of the ancestral host. In both symbiont speciation without host switching (*λ*_*S*_) and cospeciation (*λ*_*C*_), the descendant symbiont lineages retain their association with the ancestral lineage (Fig. 1), contributing to host conservatism of the symbionts. On the contrary, in symbiont speciation with host switching (*λ*_*W*_), one of the descendant symbiont lineages switches to a new host lineage, reducing the host conservatism of the symbionts (i.e., contributing to host lability). Thus, the simultaneous inference of *λ*_*S*_, *λ*_*C*_, and *λ*_*W*_ allows patterns of host conservatism to be attribute to two types of speciation increasing host conservatism and one type of speciation decreasing it. The approach presented here adds to the the existing methods to tease apart the relative contributions of different macroevolutionary processes to cophylogenetic patterns (Alcala et al., 2017; Satler et al., 2019), but under a more biologically realistic assumption that speciation and extinction occur on the timescale in the host and symbiont clades. With this improved biological realism, different speciation processes’ contributions may be more easily compared within the *same* cophylogenetic system; additionally, a certain speciation process’s contributions may also be more easily compared between *multiple* cophylogenetic systems.

In the beetle mimicry example reanalyzed in this work, we find that the two types of speciation increasing host conservatism and the one type decreasing it do not differ in rate considerably (Fig. 4). Van Dam et al. (2024) reveals that, in the beetle mimicry system, a number of interactions are conserved over long periods of time while others are more labile and transient. This mixed pattern of host conservatism, as our results suggest, could have arisen from the fact that speciation events increasing host conservatism are balanced by those decreasing it.

When a cophylogeny is seen as an evolving system of its own, questions can be asked about what affects the rates of different types of speciation in a cophylogeny. In phylogenetic comparative research, significant efforts have been devoted to studying how orgnismal traits affect speciation rates (Garamszegi, 2014; Revell and Harmon, 2022; Morlon et al., 2024). Similar questions can be asked about cophylgenetic systems. For example, key innovations have been hypothesized and debated as a driver of speciation for single clades (Miller et al., 2023) - do certain types of key innovation also affect the rates of speciation in a cophylogenetic system? In the beetle mimicry example, limited dispersal among islands has been shown to drive cospeciation Van Dam et al. (2024). Thus, one hypothesis may be that the evolution of long-distance dispersal ability may decrease the rate of cospeciation. In a different vein, the old and still influential “escape-and-radiate” hypothesis states that the evolution of novel defensive traits in symbionts allows them to “escape” from the hosts and, as a result, radiate (Ehrlich and Raven, 1964; Cogni et al., 2022). Then, what are the effects of these defensive traits on symbiont speciation without host switching, cospeciation, and symbiont speciation with host switching? Comparing the speciation rates of cophylogenetic systems with these key innovations versus those without them may provide insights into these questions and hypotheses.

An interesting potential for the approach presented here lies in the possibility of informing real-world problems about public health or conservation. For instance, the coevolutionary history of coronoviruses and their mammalian hosts has been shown to be characterized by frequent host switches (Maestri et al., 2024). Comparisons of host-switching speciation rates between different strains of viruses (which are essentially clades), enabled by the new ABC approach, would allow one to ask whether highly virulent strains switch hosts more or less frequently than less virulent strains. Alternatively, one could ask whether historical rates of host-switching speciation predict pathogens’ risk of spilling over to other host species in the future. In a different vein, specialism (i.e., high host specificity) has been hypothesized as potential evolutionary dead ends that lead to extinction (Day et al., 2016). The most extreme cases of specialism in symbionts, such as fig/fig-wasp mutualisms, tend to have arisen almost exclusively through cospeciation (Machado et al., 2005). Thus, could the degree to which cospeciation predominates in a cophylogenetic system serve as an indicator of extinction risk? Also interested in extinction risk, Mulvey et al. (2022) presents a cophylogenetic method to estimate the extinction risk of symbionts based on the number of symbiont extinctions, host-switchs, and non-host-switching events. However, this method does not take into account the speciation-extinction dynamics that occur simultaneously in the host and symbiont clades. An interesting future direction may be to develop an extinction risk index based on speciation and extinction rates estimated from a cophylogeny, such as those considered in this study.

By considering the cophylogenetic system as a whole, the new ABC approach shows the potential for some important questions in ecology and evolution to be better understood in light of rates of speciation and extinction in both the host and symbiont clades (i.e., in events per unit time). To answer these questions using the new approach, more efforts to compile time-calibrated cophylogenies will be needed.

## 5 Acknowledgments

We are grateful for the high-performance computing resources provided by the University of Arizona. We thank Kristen Martinet for feedback on the manuscript. We thank Matthew H. Van Dam for clarifying the nature of the beetle mimicry dataset. We thank members of the Data Diversity Lab for discussions about this work.

## 6 Supplementary Material

Supplementary figures and tables are available at: https://doi.org/10.6084/m9.figshare.29084336.

## 7 Data Availability

Code used in this study is available at: https://github.com/yichaozeng/cophy_ABC. Datasets used in this study are available at: https://doi.org/10.6084/m9.figshare.29084336..

## Notes

https://doi.org/10.6084/m9.figshare.29084336

### Competing Interest Statement

The authors have declared no competing interest.

