## Supplementary Material and Datasets for "Macroevolutionary Rates of Species Interactions: Approximate Bayesian Inference from Cophylogenies": Figs_S1-S5.docx

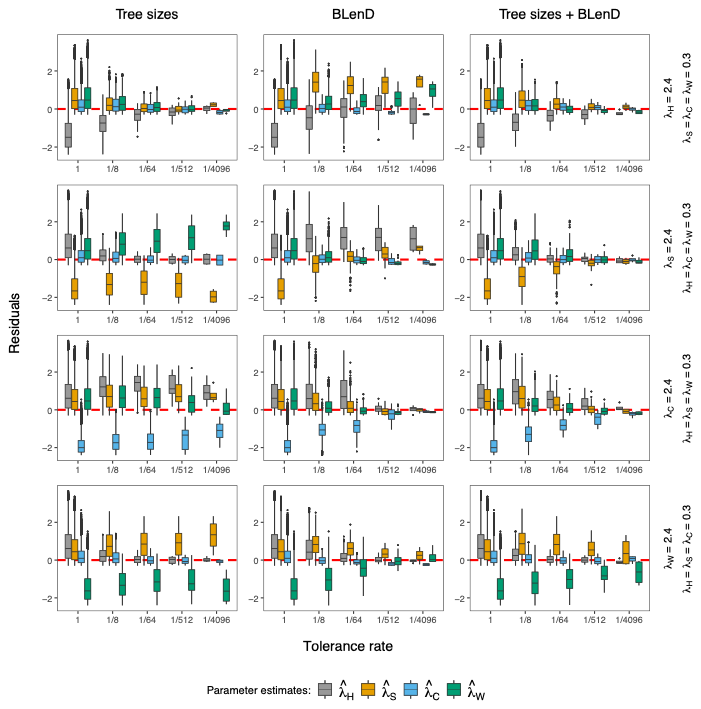

Fig. S1. Convergence plots of the speciation rate estimates $(\hat{\lambda}_{H},\hat{\lambda}_{S},\hat{\lambda}_{C},\hat{\lambda}_{W})$ when $\epsilon_{H}$ = $\epsilon_{S}$ = 0. Residuals are calculated as estimates minus the true values $(\lambda_{H},\lambda_{S},\lambda_{C},\lambda_{W})$. The estimates converge well if the residuals approach zero as tolerance rate approaches zero. Simulated cophylogenies whose true parameters are known (vertical text on the right) are used as observed data in the ABC framework. Each observed cophylogeny contains 500 replicates; the median BLenD density and tree sizes are taken across all replicates before being used as observations in the ABC.

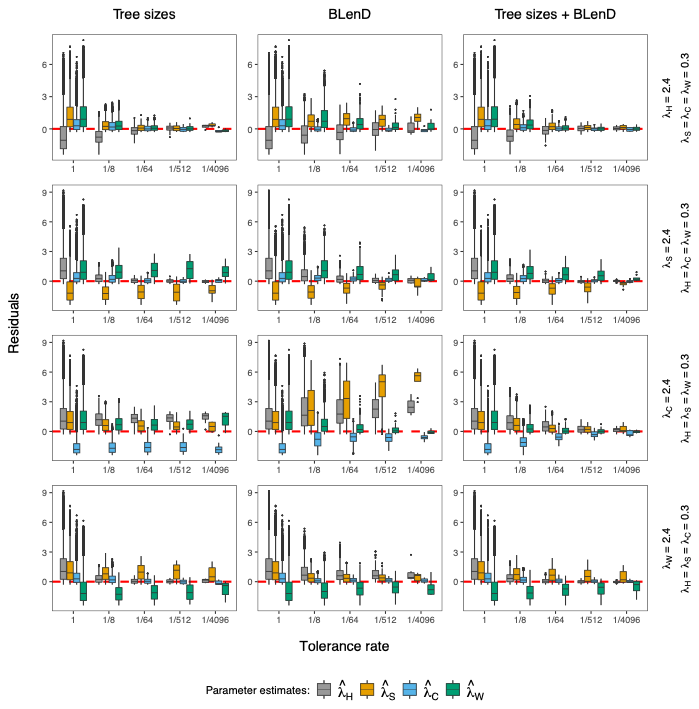

Fig. S2. Convergence plots of the speciation rate estimates $(\hat{\lambda}_{H},\hat{\lambda}_{S},\hat{\lambda}_{C},\hat{\lambda}_{W})$ when $\epsilon_{H}$ = 0.7, $\epsilon_{S}$ = 0. Residuals are calculated as estimates minus the true values $(\lambda_{H},\lambda_{S},\lambda_{C},\lambda_{W})$. The estimates converge well if the residuals approach zero as tolerance rate approaches zero. Simulated cophylogenies whose true parameters are known (vertical text on the right) are used as observed data in the ABC framework. Each observed cophylogeny contains 500 replicates; the median BLenD density and tree sizes are taken across all replicates before being used as observations in the ABC.

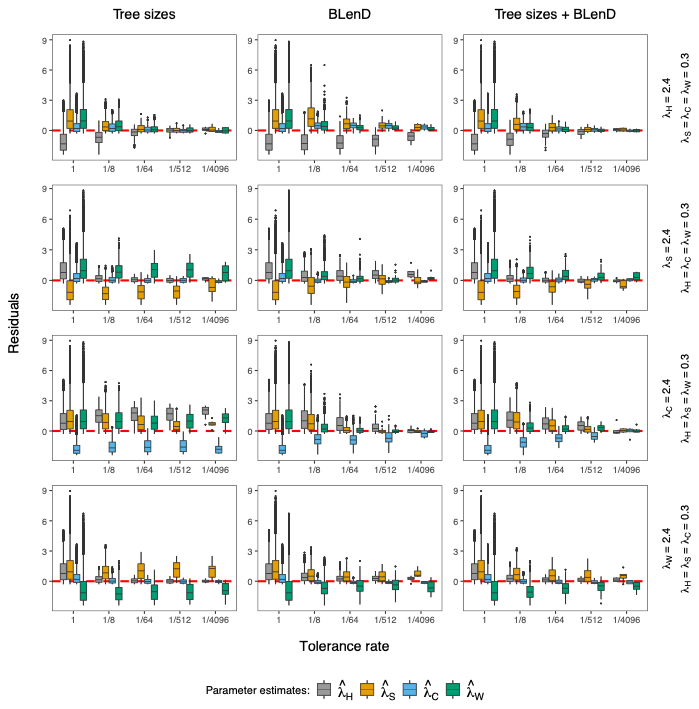

Fig. S3. Convergence plots of the speciation rate estimates $(\hat{\lambda}_{H},\hat{\lambda}_{S},\hat{\lambda}_{C},\hat{\lambda}_{W})$ when $\epsilon_{H}$ = 0, $\epsilon_{S}$ = 0.7. Residuals are calculated as estimates minus the true values $(\lambda_{H},\lambda_{S},\lambda_{C},\lambda_{W})$. The estimates converge well if the residuals approach zero as tolerance rate approaches zero. Simulated cophylogenies whose true parameters are known (vertical text on the right) are used as observed data in the ABC framework. Each observed cophylogeny contains 500 replicates; the median BLenD density and tree sizes are taken across all replicates before being used as observations in the ABC.

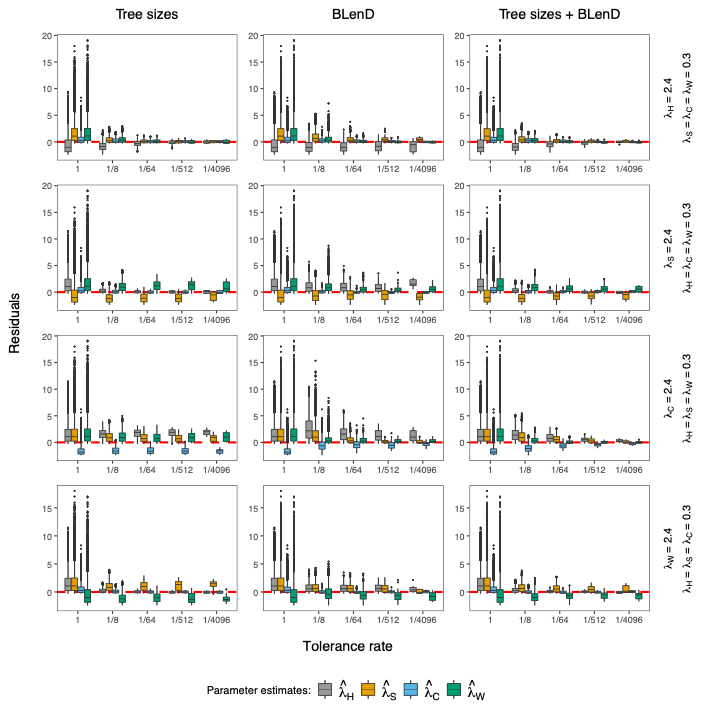

Fig. S4. Convergence plots of the speciation rate estimates $(\hat{\lambda}_{H},\hat{\lambda}_{S},\hat{\lambda}_{C},\hat{\lambda}_{W})$ when ${\epsilon_{H}= \epsilon}_{S}$ = 0.7. Residuals are calculated as estimates minus the true values $(\lambda_{H},\lambda_{S},\lambda_{C},\lambda_{W})$. The estimates converge well if the residuals approach zero as tolerance rate approaches zero. Simulated cophylogenies whose true parameters are known (vertical text on the right) are used as observed data in the ABC framework. Each observed cophylogeny contains 500 replicates; the median BLenD density and tree sizes are taken across all replicates before being used as observations in the ABC.

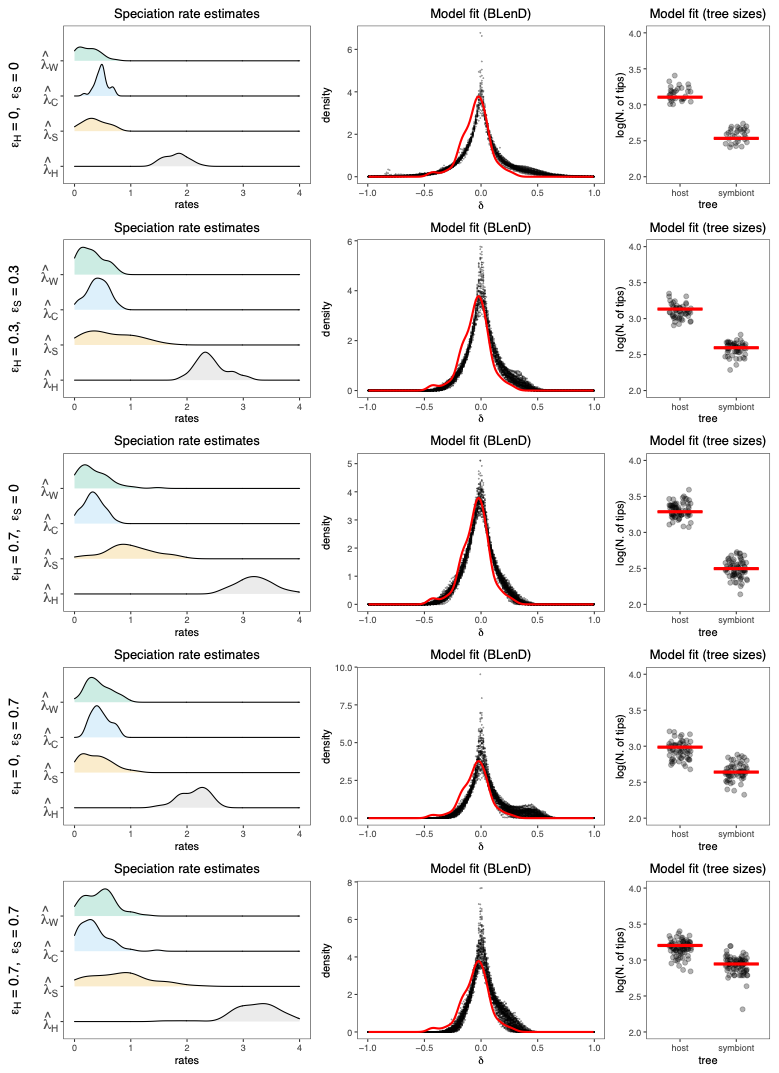

Figure 4. Speciation rate inference from the cophylogenetic dataset of beetle mimicry (Van Dam et al. 2024) under different assumptions of $\epsilon_{H}$ and $\epsilon_{S}$. Four rate parameters are considered: $\lambda_{H}$- host speciation; $\lambda_{S}$- host speciation without host switching; $\lambda_{C}$ - cospeciation; $\lambda_{W}$ - symbiont speciation with host switching. Both the BLenD curve and tree sizes are used in tandem as summary statistics. The tolerance rate used is 1/512. **Left**: Density curves of the speciation rate estimates. **Middle & Right**: Posterior predictive checks of model fit. Shown here are the BLenD curve and tree sizes of the beetle data (solid line) and accepted simulations in the ABC (dots).
