## Supplementary Material and Datasets for "Macroevolutionary Rates of Species Interactions: Approximate Bayesian Inference from Cophylogenies": Supplementary_Information.docx

The paper contains the following as supplementary information and datasets:

- *Figs_S1-S5.docx* – Figs. S1-S5
- *Tables_S1-S2.docx* – Tables S1-S2
- Datasets:

1. *para_est_*[$\varepsilon_{H}$]*_*[$\varepsilon_{S}$]*.rds –* 5 files corresponding to different assumptions of $\varepsilon_{H}$ and $\varepsilon_{S}$. Each file contains 12 components that contain the parameter estimates used to create the convergence plots (Figs. 3, S1-S4). Each panel in a convergence plot (Figs. 3, S1-S4) corresponds to one of the 12 components as follows:

*1st	5th	9th
2nd	6th	10th
3rd	7th	11th
4th	8th	12th*

2. *cophy_real.rds* – The beetle mimicry cophylogeny from (Van Dam et al. 2024). This dataset has been rescaled such that: (i) both the host and symbiont phylogenies contain a root edge (branch); (ii) both the host and symbiont phylogenies are of the same height (from the root to the tips, including the length of the root edge); (iii) both the host and symbiont phylogenies are of the same height as the cophylogenies simulated with treeducken (2 unit time in this case).
3. *real_para_est.rds* – Speciation rate estimates from the beetle mimicry dataset (Van Dam et al. 2024). This R dataset contains 5 components, each corresponding to results obtained under a different assumption of $\varepsilon_{H}$ and $\varepsilon_{S}$ (0/0, 0.3/0.3, 0.7/0, 0/0.7, 0.7/0.7 for $\varepsilon_{H}$ and $\varepsilon_{S}$, respectively). Note that $\lambda_{W}$ is represented by exp_H (consistent with the original treeducken denotation in Dismukes & Heath 2022) in this R dataset.
