## Supplementary Material and Datasets for "Macroevolutionary Rates of Species Interactions: Approximate Bayesian Inference from Cophylogenies": Tables_S1-S2.docx

Table S1. Performance of speciation rate estimation and speciation rate heterogeneity detection. Only the tree sizes are used as summary statistics. The tolerance rate used is 1/4096.

| Relative extinction rates | | True speciation rates | | | | Rate estimates | | | | Rate heterogeneity detection | |
| --- | --- | --- | --- | --- | --- | --- | --- | --- | --- | --- | --- |
| $\epsilon_{H}$ | $\epsilon_{S}$ | $\lambda_{H}$ | $\lambda_{S}$ | $\lambda_{C}$ | $\lambda_{W}$ | $\hat{\lambda}_{H}$ | $\hat{\lambda}_{S}$ | $\hat{\lambda}_{C}$ | $\hat{\lambda}_{W}$ | Correctness | Lift |
| 0 | 0 | **2.4** | 0.3 | 0.3 | 0.3 | **2.4 (0.2)** | 0.52 (0.11) | 0.13 (0.13) | 0.2 (0.09) | 0.994 | 3.976 |
| 0 | 0 | 0.3 | **2.4** | 0.3 | 0.3 | 0.32 (0.3) | **0.45 (0.36)** | 0.27 (0.3) | 2.08 (0.49) | 0.476 | 1.904 |
| 0 | 0 | 0.3 | 0.3 | **2.4** | 0.3 | 1.25 (0.7) | 1.08 (0.45) | **1.26 (0.69)** | 0.48 (0.65) | 0.1 | 0.4 |
| 0 | 0 | 0.3 | 0.3 | 0.3 | **2.4** | 0.33 (0.13) | 1.61 (0.91) | 0.21 (0.14) | **0.87 (0.85)** | 0.576 | 2.304 |
| 0.3 | 0.3 | **2.4** | 0.3 | 0.3 | 0.3 | **2.19 (0.35)** | 0.21 (0.2) | 0.46 (0.22) | 0.25 (0.14) | 0.996 | 3.984 |
| 0.3 | 0.3 | 0.3 | **2.4** | 0.3 | 0.3 | 0.5 (0.1) | **1.86 (0.84)** | 0.11 (0.09) | 0.96 (0.85) | 0.434 | 1.736 |
| 0.3 | 0.3 | 0.3 | 0.3 | **2.4** | 0.3 | 2.11 (0.5) | 1.37 (0.82) | **0.59 (0.54)** | 1.06 (0.92) | 0.078 | 0.312 |
| 0.3 | 0.3 | 0.3 | 0.3 | 0.3 | **2.4** | 0.25 (0.16) | 1.55 (0.66) | 0.38 (0.16) | **1.15 (0.71)** | 0.528 | 2.112 |
| 0.7 | 0 | **2.4** | 0.3 | 0.3 | 0.3 | **2.62 (0.14)** | 0.65 (0.15) | 0.11 (0.14) | 0.14 (0.12) | 0.998 | 3.992 |
| 0.7 | 0 | 0.3 | **2.4** | 0.3 | 0.3 | 0.24 (0.15) | **1.5 (0.81)** | 0.36 (0.2) | 1.25 (0.87) | 0.564 | 2.256 |
| 0.7 | 0 | 0.3 | 0.3 | **2.4** | 0.3 | 1.7 (0.57) | 0.72 (0.45) | **0.68 (0.61)** | 1.41 (0.86) | 0.042 | 0.168 |
| 0.7 | 0 | 0.3 | 0.3 | 0.3 | **2.4** | 0.39 (0.16) | 1.04 (0.77) | 0.2 (0.19) | **1.64 (0.78)** | 0.546 | 2.184 |
| 0 | 0.7 | **2.4** | 0.3 | 0.3 | 0.3 | **2.51 (0.21)** | 0.44 (0.28) | 0.23 (0.16) | 0.3 (0.27) | 0.996 | 3.984 |
| 0 | 0.7 | 0.3 | **2.4** | 0.3 | 0.3 | 0.42 (0.16) | **1.76 (0.88)** | 0.19 (0.16) | 1.05 (0.81) | 0.48 | 1.92 |
| 0 | 0.7 | 0.3 | 0.3 | **2.4** | 0.3 | 2.19 (0.67) | 1 (0.34) | **0.7 (0.59)** | 1.43 (0.89) | 0.058 | 0.232 |
| 0 | 0.7 | 0.3 | 0.3 | 0.3 | **2.4** | 0.32 (0.14) | 1.39 (0.84) | 0.25 (0.16) | **1.52 (0.8)** | 0.548 | 2.192 |
| 0.7 | 0.7 | **2.4** | 0.3 | 0.3 | 0.3 | **2.39 (0.29)** | 0.32 (0.2) | 0.31 (0.17) | 0.33 (0.31) | 0.998 | 3.992 |
| 0.7 | 0.7 | 0.3 | **2.4** | 0.3 | 0.3 | 0.35 (0.26) | **1.66 (0.84)** | 0.32 (0.27) | 1.27 (1.06) | 0.514 | 2.056 |
| 0.7 | 0.7 | 0.3 | 0.3 | **2.4** | 0.3 | 2.2 (0.56) | 1.16 (0.75) | **0.73 (0.46)** | 1.35 (0.88) | 0.006 | 0.024 |
| 0.7 | 0.7 | 0.3 | 0.3 | 0.3 | **2.4** | 0.27 (0.17) | 1.58 (0.83) | 0.27 (0.19) | **1.22 (0.81)** | 0.526 | 2.104 |

Table S2. Performance of speciation rate estimation and speciation rate heterogeneity detection. Only the BLenD curve is used in tandem as summary statistics. The tolerance rate used is 1/4096.

| Relative extinction rates | | True speciation rates | | | | Rate estimates | | | | Rate heterogeneity detection | |
| --- | --- | --- | --- | --- | --- | --- | --- | --- | --- | --- | --- |
| $\epsilon_{H}$ | $\epsilon_{S}$ | $\lambda_{H}$ | $\lambda_{S}$ | $\lambda_{C}$ | $\lambda_{W}$ | $\hat{\lambda}_{H}$ | $\hat{\lambda}_{S}$ | $\hat{\lambda}_{C}$ | $\hat{\lambda}_{W}$ | Correctness | Lift |
| 0 | 0 | **2.4** | 0.3 | 0.3 | 0.3 | **2.22 (1.1)** | 1.58 (0.73) | 0.02 (0.02) | 1.2 (0.61) | 0.604 | 2.416 |
| 0 | 0 | 0.3 | **2.4** | 0.3 | 0.3 | 1.41 (0.74) | **2.96 (0.2)** | 0.16 (0.13) | 0.04 (0.04) | 0.418 | 1.672 |
| 0 | 0 | 0.3 | 0.3 | **2.4** | 0.3 | 0.32 (0.21) | 0.27 (0.11) | **2.31 (0.07)** | 0.17 (0.02) | 0.868 | 3.472 |
| 0 | 0 | 0.3 | 0.3 | 0.3 | **2.4** | 0.25 (0.08) | 0.54 (0.38) | 0.07 (0.05) | **2.58 (0.41)** | 0.678 | 2.712 |
| 0.3 | 0.3 | **2.4** | 0.3 | 0.3 | 0.3 | **2.23 (0.76)** | 0.91 (0.33) | 0.52 (0.27) | 0.55 (0.17) | 0.462 | 1.848 |
| 0.3 | 0.3 | 0.3 | **2.4** | 0.3 | 0.3 | 0.76 (0.57) | **2.4 (0.49)** | 0.16 (0.12) | 0.64 (0.23) | 0.24 | 0.96 |
| 0.3 | 0.3 | 0.3 | 0.3 | **2.4** | 0.3 | 0.43 (0.38) | 0.36 (0.18) | **2.51 (0.39)** | 0.38 (0.18) | 0.564 | 2.256 |
| 0.3 | 0.3 | 0.3 | 0.3 | 0.3 | **2.4** | 0.45 (0.31) | 0.49 (0.29) | 0.23 (0.08) | **2.27 (0.55)** | 0.496 | 1.984 |
| 0.7 | 0 | **2.4** | 0.3 | 0.3 | 0.3 | **2.53 (0.6)** | 1.32 (0.68) | 0.18 (0.17) | 0.61 (0.66) | 0.52 | 2.08 |
| 0.7 | 0 | 0.3 | **2.4** | 0.3 | 0.3 | 0.39 (0.29) | **2.09 (0.63)** | 0.42 (0.16) | 0.74 (0.5) | 0.268 | 1.072 |
| 0.7 | 0 | 0.3 | 0.3 | **2.4** | 0.3 | 2.81 (0.72) | 5.5 (1.23) | **1.79 (0.33)** | 0.23 (0.18) | 0.432 | 1.728 |
| 0.7 | 0 | 0.3 | 0.3 | 0.3 | **2.4** | 1.07 (0.78) | 0.74 (0.43) | 0.41 (0.22) | **1.78 (0.78)** | 0.388 | 1.552 |
| 0 | 0.7 | **2.4** | 0.3 | 0.3 | 0.3 | **1.77 (0.57)** | 0.63 (0.29) | 0.65 (0.25) | 0.51 (0.24) | 0.438 | 1.752 |
| 0 | 0.7 | 0.3 | **2.4** | 0.3 | 0.3 | 0.99 (0.49) | **2.39 (0.56)** | 0.21 (0.07) | 0.56 (0.3) | 0.242 | 0.968 |
| 0 | 0.7 | 0.3 | 0.3 | **2.4** | 0.3 | 0.34 (0.25) | 0.28 (0.14) | **2.14 (0.37)** | 0.24 (0.11) | 0.624 | 2.496 |
| 0 | 0.7 | 0.3 | 0.3 | 0.3 | **2.4** | 0.54 (0.24) | 1.08 (0.38) | 0.17 (0.11) | **1.81 (0.66)** | 0.604 | 2.416 |
| 0.7 | 0.7 | **2.4** | 0.3 | 0.3 | 0.3 | **1.59 (1.1)** | 0.75 (0.38) | 0.3 (0.13) | 0.26 (0.18) | 0.41 | 1.64 |
| 0.7 | 0.7 | 0.3 | **2.4** | 0.3 | 0.3 | 1.99 (0.72) | **1.49 (0.8)** | 0.25 (0.11) | 0.97 (0.83) | 0.162 | 0.648 |
| 0.7 | 0.7 | 0.3 | 0.3 | **2.4** | 0.3 | 1.53 (1.04) | 0.51 (0.47) | **2.13 (0.59)** | 0.62 (0.46) | 0.278 | 1.112 |
| 0.7 | 0.7 | 0.3 | 0.3 | 0.3 | **2.4** | 0.84 (0.67) | 0.38 (0.34) | 0.37 (0.22) | **1.62 (0.78)** | 0.518 | 2.072 |
